# Complement Regulator Factor H is a Cofactor for Thrombin in both Pro- and Anticoagulant Roles

**DOI:** 10.1101/2021.07.22.452893

**Authors:** Genevieve. McCluskey, Gemma E. Davies, Rebekah L. Velounias, Tim R. Hughes, B. Paul Morgan, Roger J. S. Preston, Peter W. Collins, P. Vincent Jenkins, Meike Heurich

**Affiliations:** School of Pharmacy and Pharmaceutical Sciences, College of Biomedical and Life Sciences, Cardiff University, United Kingdom; School of Medicine, College of Biomedical and Life Sciences, Cardiff University, United Kingdom; Department of Haematology, University Hospital of Wales, Cardiff and Vale University Health Board, United Kingdom; Cardiff Haemophilia Centre, University Hospital of Wales, Cardiff and Vale University Health Board, United Kingdom; Irish Centre for Vascular Biology, School of Pharmacy & Biomolecular Sciences, Royal College of Surgeons in Ireland, Ireland

**Keywords:** Complement, coagulation, factor H, thrombin, fibrin, activated protein C

## Abstract

**Background:** Complement FH (FH) is a key regulator of complement activity whereas thrombin (FIIa) is central to hemostasis with both pro- and anticoagulant functions. Both have separately been shown to have auxiliary activities across the two systems. The purpose of this study was to determine the effect of FH on pro- and anti-coagulant functions and investigate the interaction between FH and thrombin.

**Methods:** Tail bleeding time and hemolysis were measured in FH-deficient mice (*CFH*^−/−^). Activated partial thromboplastin time (aPTT) was determined in FH-depleted human plasma. FH effect on fibrin clot generation was investigated in turbidity assays and on activated protein C (APC) generation. Binding affinity of thrombin with FH was determined using surface plasmon resonance (SPR).

**Results:** Tail bleeding time in *CFH*^−/−^ mice was significantly prolonged compared to wild type mice. The aPTT in FH-depleted human plasma was elevated compared to normal plasma and restored by adding back FH to depleted plasma. Accordingly, FH enhanced thrombin-mediated fibrin clot generation by shortening lag time, increasing rate of clot formation and maximum turbidity, and affected clot structure. Despite this, FH also increased the rate of thrombin-mediated protein C (PC) activation, both in the presence and absence of soluble recombinant thrombomodulin (TM). Nanomolar affinity binding of FH with thrombin, but not prothrombin, was confirmed.

**Conclusion:** Complement FH binds thrombin with strong affinity and acts as a novel cofactor that enhances both pro- and anticoagulant actions of thrombin. These data highlight an important role for FH in hemostasis.

**Key points:**

- Absence of FH prolongs tail bleeding time in *CFH*^−/−^ mice and activated partial thromboplastin time (aPTT) is elevated in human FH-depleted plasma.
- FH acts a cofactor for thrombin by enhancing fibrin generation, altering fibrin clot structure and enhancing TM-thrombin mediated protein C activation

## Introduction

The blood-borne coagulation and complement systems share evolutionary origins with and molecular crosstalk (1–3). Central to coagulation is serine protease thrombin, mediating both pro- and anticoagulant functions (4–6). Thrombin has numerous substrates (6,7) and diverse roles (8–10); cofactors modulate thrombin substrate recognition in the hemostatic response to vascular injury (6,7). Thrombin acts predominantly as a procoagulant by cleaving fibrinogen to fibrin (12), with its structure and properties affected by endogenous mediators (12,13) and thrombin concentration (12,14).

Thrombin specificity for procoagulant substrates is altered by binding to its cofactor thrombomodulin (TM) (15), enabling anticoagulant activated protein C (APC) generation (16,17). TM is expressed and best characterized on endothelial cells (18,19), where it also exerts non-hemostatic functions (20) including the enhancement of complement regulator FH-mediated regulation of complement (21,22).

The complement system, integral to non-cellular innate immunity, comprises blood and membrane-bound proteins generating opsonins (C3b), anaphylatoxins (C3a, C5a) and the membrane attack complex (24) in response to vascular injury (25) and infection (26). Complement is regulated on self-cells (27) with factor H (FH) as the key regulator of the complement alternative pathway (28), acting as C3 convertase decay accelerator and cofactor for serine protease factor I (FI) mediated inactivation of C3b (29–31).

FH deficiency results in uncontrolled alternative pathway activation and secondary C3 depletion (32). FH deficiency is associated with renal pathologies membranoproliferative glomerulonephritis II (MPGNII) (33) or atypical hemolytic uremic syndrome (aHUS) (34) - a thrombotic microangiopathy (TMA) characterized by microvascular occlusion through platelet thrombi, thrombocytopenia, and microangiopathic hemolytic anemia (35–37). Complement deficiency knockout mouse models lacking C3 (38,39), C5 (40), C3aR (41) or C6 (42) all affect hemostasis; showing altered bleeding and thrombus formation.

There are numerous points of intersection between complement and coagulation pathways that may cause a prothrombotic phenotype in disease. For instance, both FH and thrombin have auxiliary activities across the pathways. Thrombin is able to cleave C3 and C5 to generate C3a, C5a (43) and C5b-9 (46), which could directly induce tissue factor (TF) expression (45,46). FH is a fibrin clot component (47,48), and has several coagulation protein ligands, including fibrin(ogen) (49), and FXII (50,51), von Willebrand factor (52,53) and platelets (54–56). However, no direct physiologic function of FH in the coagulation system has been previously demonstrated. Our study aimed to establish the role of FH in hemostatic, pro- and anticoagulant mechanisms.

## Methods

### Animal studies

Animal work was carried out at Cardiff University under the authority of UK Home Office license PPL 30/3365 and complied with ARRIVE guidelines. In brief, 20 week old, male, wild-type (WT) or FH-deficient (*CFH*^−/−^) mice on a C57/Bl6 background, generated as described previously (57), were anaesthetized and a 3-mm portion of distal tail tip was amputated. Tail bleeding time was monitored and experiments stopped at 20 minutes (**Supplementary**). Whole blood was obtained by cardiac puncture, and harvested into EDTA and used to measure plasma hemoglobin (plasma-Hb) and quantify soluble recombinant thrombomodulin (sTM-ELISA, ab209880, Abcam) and thrombin-antithrombin (TAT-ELISA, ab137994, Abcam).

### FH purification and FH-depletion from plasma

Pooled human citrated or EDTA-plasma was obtained from consenting, healthy donors. Citrated plasma was immunoaffinity-depleted of factor B (FB) and FH by flowing over anti-FB monoclonal antibody (mAb) (JC1; to prevent activation of the alternative complement pathway and complement consumption) and anti-FH mAb (35H9) on Hi-Trap HP-NHS columns (GE Healthcare). FH protein was purified from EDTA-plasma using an anti-FH mAb (35H9) immunoaffinity column. Affinity depletion of FH and FB in citrated plasma was verified by Western Blot and FH depletion confirmed by ELISA (HK342-02, Hycylt, NL) and residual alternative pathway activation was determined by visualizing FB activation fragment Bb using SDS-PAGE and anti-FB/Bb (JC1) mAb on Western blot (**Supplementary Figure 1**).

### Activated partial thromboplastin time (aPTT) and partial thromboplastin time (PT) analysis

Activated partial thromboplastin time (aPTT) and prothrombin time (PT) were determined in human citrated depleted plasma (ΔFB/FH or ΔFH) compared to normal plasma. Citrated ΔFB/FH or ΔFH-plasma (50μl), with restored 1μM FH (or buffer control), was mixed with 50μl HemosIL SynthASil aPTT reagent (120 seconds) and coagulation initiated by addition of 50μl of 20mM CaCl_2_; PT was performed by addition of 50ul HemosIL RecombiPlastin 2G PT reagent to plasma; and measured by ACL TOP 750 (Werfen, Warrington, UK).

### Fibrin clot turbidity

Thrombin (2.5nM, Enzyme Research Laboratories, Swansea, UK) in the presence or absence of FH (1.5-100nM) was added to 4g/L (11.8μM) plasminogen-depleted fibrinogen (Enzyme Research Laboratories, UK) in 10mM HEPES, 150 mM NaCl, 5 mM CaCl_2_, buffer pH7.4 and fibrin clot generation was monitored by assessing turbidity at A405nm every 60 seconds for 120 minutes.

### Fluorescent microscopy

Fibrin clots were prepared as described for the turbidity assays with the following modification. Clots were formed in chambered coverslips within an area of 1cm^2^ defined with an ImmunoPen (402176, Merck Millipore). AF488-conjugated fibrinogen was added (fibrinogen-AF, Invitrogen, St Louis, MO, USA) at 1:10 (v/v) ratio to 11.8μM plasminogen-free fibrinogen (Enzyme Research Laboratories, UK) prior to 2.5nM thrombin and 100nM FH in 10mM HEPES, 150mM NaCl, 5mM CaCl_2_, pH7.4 or buffer control and left to mature for 2◻hours at 37◻°C. Images were acquired by fluorescence microscopy (EVOS™ M7000 Imaging System, ThermoFisher, UK) at 20X magnification. ImageJ Version 1.43 (U.S. National Institutes of Health, USA) was used to determine fiber density by counting the number of fibers crossing 25 lines of 100μm spacing placed in the image (25 lines per image, 12 images taken in different areas of the clot) using the plug-in grid. Each experimental condition was repeated 10 times.

### Activated protein C generation assay

The effect of FH on protein C (PC) activation was monitored by combining FH (0.5-125nM) with 1.75nM thrombin and 200nM PC (Haematologic Technologies, Inc., USA) in the absence or presence of 10nM soluble recombinant TM (R&D Systems/Biotechne, USA). Activation of PC (6.25-800nM) was determined by addition of 1.75nM thrombin in the presence or absence of 100nM FH and 10nM TM; incubating for 2 hours at 37 ◻C, followed by 1U/well Hirudin (Sigma, UK) for 10 minutes. APC substrate (1.25mg/ml BIOPHEN™ CS-21(66), Hyphen Biomed, USA) was added and absorbance read at A405nm for 1 hour at 60 second intervals.

### In vitro complex formation between FH and thrombin

Direct binding of FH with thrombin was determined by adsorption of 10μg/ml PPACK-thrombin or prothrombin (Enzyme Research Laboratories, UK), in carbonate buffer, pH9.6, onto 96-well microtiter plates overnight at 4°C, followed by blocking (2.5% milk for 2 hours at 37°C) and washing in PBST. FH (0.5-32nM) was incubated for 2 hours at 37°C in the wells, unbound material washed off, and bound FH detected using HRP-conjugated polyclonal rabbit anti-FH antibody (1μg/ml; in-house). Substrate (OPD, O-phenylenediamine dihydrochloride, Thermo Scientific) was added, the reaction stopped using 10% H_2_SO_4_ and read at A492nm.

An indirect capture ELISA for thrombin-FH complexes was developed by capturing FH (20μg/ml) via adsorbed anti-FH mAb (10μg/ml in carbonate buffer pH9.6 for 24 hours at 4°C, followed by blocking in 2.5% (w/v) milk/PBST and washing in PBST. PPACK-thrombin titration series (17-540nM) was added for 1 hour at room temperature, and unbound material washed off in PBST. FH-thrombin complex was detected by adding 1μg/ml polyclonal sheep anti-thrombin antibody (ab61367, Abcam) in 5% (w/v) milk PBST for 1hour at 37°C, followed by wash and incubation with 0.5μg/ml secondary HRP-conjugated anti-sheep antibody (Jackson ImmunoResearch, USA) before washing in PBST and addition of substrate (OPD, O-phenylenediamine dihydrochloride, Thermo Scientific). The reaction was stopped using 10% H_2_SO_4_ and read at A492nm.The formation of FH-thrombin complex in normal human plasma was measured by adding 1μM PPACK-thrombin to 50μl EDTA-plasma containing endogenous FH and quantified using a FH-thrombin standard curve.

### SPR analysis of FH with thrombin

SPR analysis was performed on a Biacore T200 instrument on a Series S CM5 chip (GE Healthcare) in 10mM HEPES, 150mM NaCl, 0.001% P20, pH7.4. PPACK-thrombin or fibrinogen in acetate buffer pH4.5 was immobilized to a chip surface (1000 or 10000 RU) using NHS/EDC-amine coupling and residual binding sites blocked by 1M ethanolamine (GE Healthcare). FH (0.5-62nM) was flowed over immobilized thrombin or fibrinogen and affinity (KD) was analyzed using BiaEvaluation software version 2.0.

### Complement hemolysis assay

Thrombin (500nM) or buffer control is added to 10% normal human serum (NHS), followed by serial dilution in complement fixing buffer (CFD, Oxoid, UK) and addition to 2% Amboceptor-sensitised sheep erythrocytes (ShEA). Alternatively, a FH titration (0.49-500nM) is added to 0.625% NHS with added thrombin or PPACK-thrombin (10nM, 500nM) or buffer control, followed by addition to 2% ShEA CFD, followed by incubation at 37 °C for one hour and centrifugation at 1500rpm for 5 minutes. Supernatant hemoglobin is measured at A405nm and percentage hemolysis calculated.

### Statistical analysis

All analyses were carried out using GraphPad Prism version 8.0.0 for Windows, GraphPad Software, San Diego, California USA. Nonparametric Mann-Whitney test or Spearman correlation was used to assess associations between concentration variables. Values of p◻<◻0.05 were considered statistically significant. All experiments were repeated at least three times.

### Data sharing statement

For original data, please contact: Meike Heurich, School of Pharmacy and Pharmaceutical Sciences, Cardiff University, United Kingdom;

## Results

Given that thrombin is a structural homologue of the complement protease FI (30), and that FH is a key cofactor for FI activity, we hypothesized that FH may affect coagulation directly by regulating thrombin activity.

### Absence of FH increases bleeding time and hemostatic ex-vivo parameters in FH-deficient (*CFH*^−/−^) mice

We hypothesized that absence of FH would lead to coagulation dysregulation in vivo. To test this, we monitored tail bleeding time and hemolysis in *CFH*^−/−^ mice, and quantified thrombin-antithrombin complexes and soluble TM levels in mouse plasma. (**Figure 1**).

**Figure 1.**
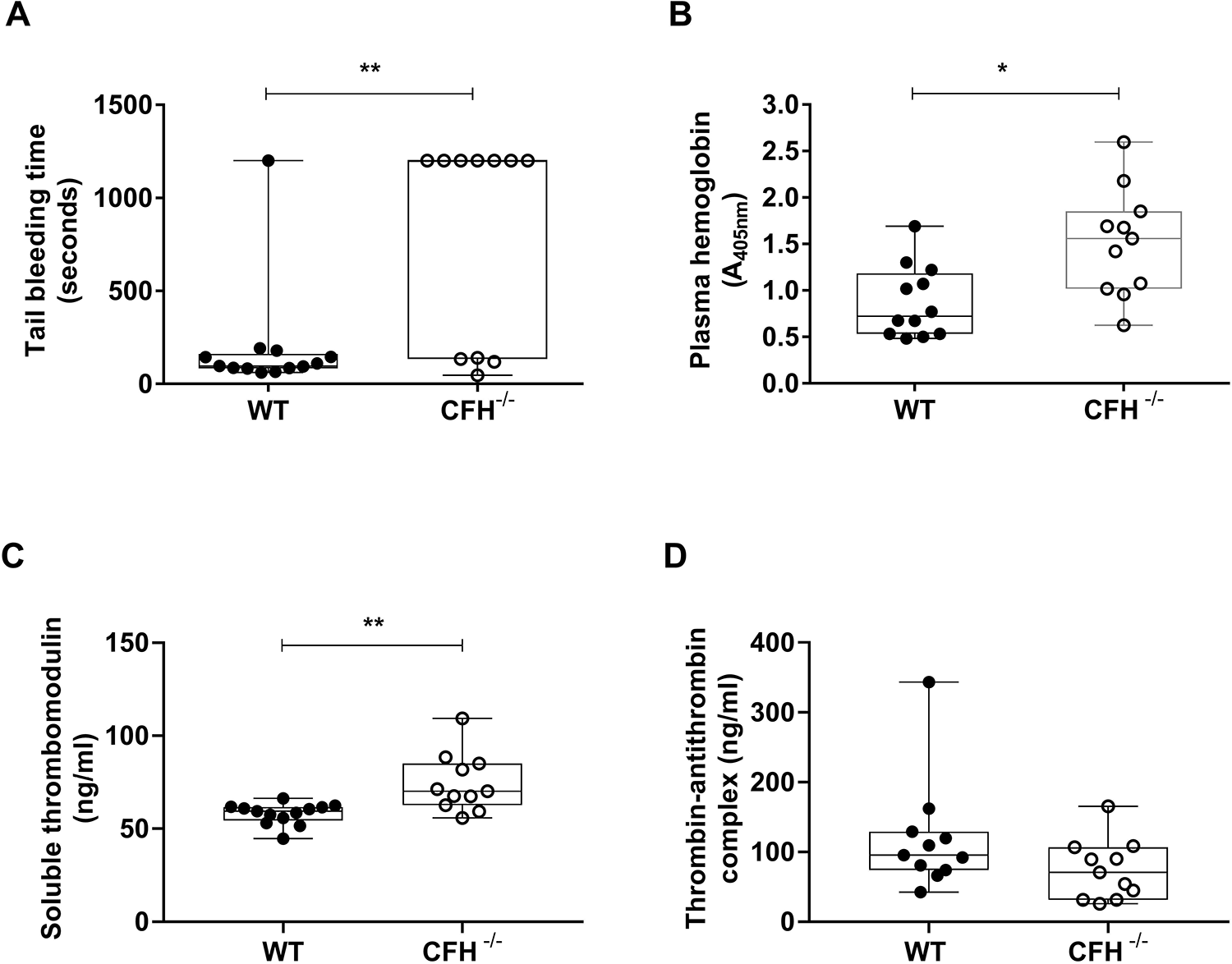
Bleeding time is prolonged in *CFH*^−/−^ mice. Tests were performed blind to genotype, in male mice, age 20 weeks. Tail bleeding assay was performed by 3mm-tail tip amputation under general anesthesia, immersing the tail in PBS at 37°C and monitoring time (seconds) until cessation of bleeding or end of experiment at 20 minutes. **(A)** *CFH^−/−^* (○) mice showed prolonged total bleeding time compared to WT (*CFH^−/−^* median: 1200 (47-1200) seconds, 25% percentile: 134, 75% percentile 1200, n=11, P=0.005; wild-type (WT, •); median: 97 (62-1200) seconds, 25% percentile: 84, 75% percentile 162, n=13. Not normally distributed bi-modal data (D’Agostino & Pearson normality test), data is expressed as median and percentile range (minimum to maximum, and percentile range) and P value calculated using Welch two-sample t test. **(B)** Plasma hemoglobin (plasma-Hb) was increased in *CFH^−/−^* plasma (1.51±0.57) compared to WT (0.87±0.39, P=0.009). Ex-vivo hemostatic parameters were quantified using standardized ELISA. **(C)** Soluble TM was significantly increased in *CFH^−/−^* plasma (74.44±15.56ng/ml, P=0.0012) compared to WT (58.03±5.6ng/ml). **(D)** Thrombin-antithrombin complex, was slightly reduced in *CFH^−/−^* plasma compared to WT (74.49±42.77ng/ml vs. 119.5±81.14ng/ml, P=0.0879), but the difference was not statistically significant. Data expressed as mean±SD and analyzed using non-parametric Mann–Whitney test.

*CFH^−/−^* mice showed significantly prolonged total tail bleeding time (**Figure 1A**), demonstrating an in vivo procoagulant role for complement FH. Seven of 11 *CFH^−/−^* mice continued to bleed through to the experiment endpoint at 1200 seconds with a median bleeding time of 1200 seconds (s) (47-1200s) compared to wildtype (median: 97s (62-1200s), P=0.017). Elevated plasma hemoglobin in *CFH^−/−^* mice, but not wildtype, suggests increased intravascular hemolysis (**Figure 1B**) perhaps induced by complement activation as observed in humans with FH deficiency (58–60). Significantly increased soluble TM was observed in *CFH^−/−^* mice (**Figure 1C**). We observed no change in thrombin-antithrombin complex indicating no significant impact on thrombin production and activation of the coagulation cascade in *CFH^−/−^* mice (**Figure 1D**).

### Activated partial thromboplastin time (aPTT) is increased in FH-depleted human plasma

To investigate the hemostatic role of FH in human plasma, we measured activated partial thromboplastin time (aPTT) and prothrombin time (PT) in FH-depleted human plasma with or without restoration of FH levels (**Figure 2**).

**Figure 2.**
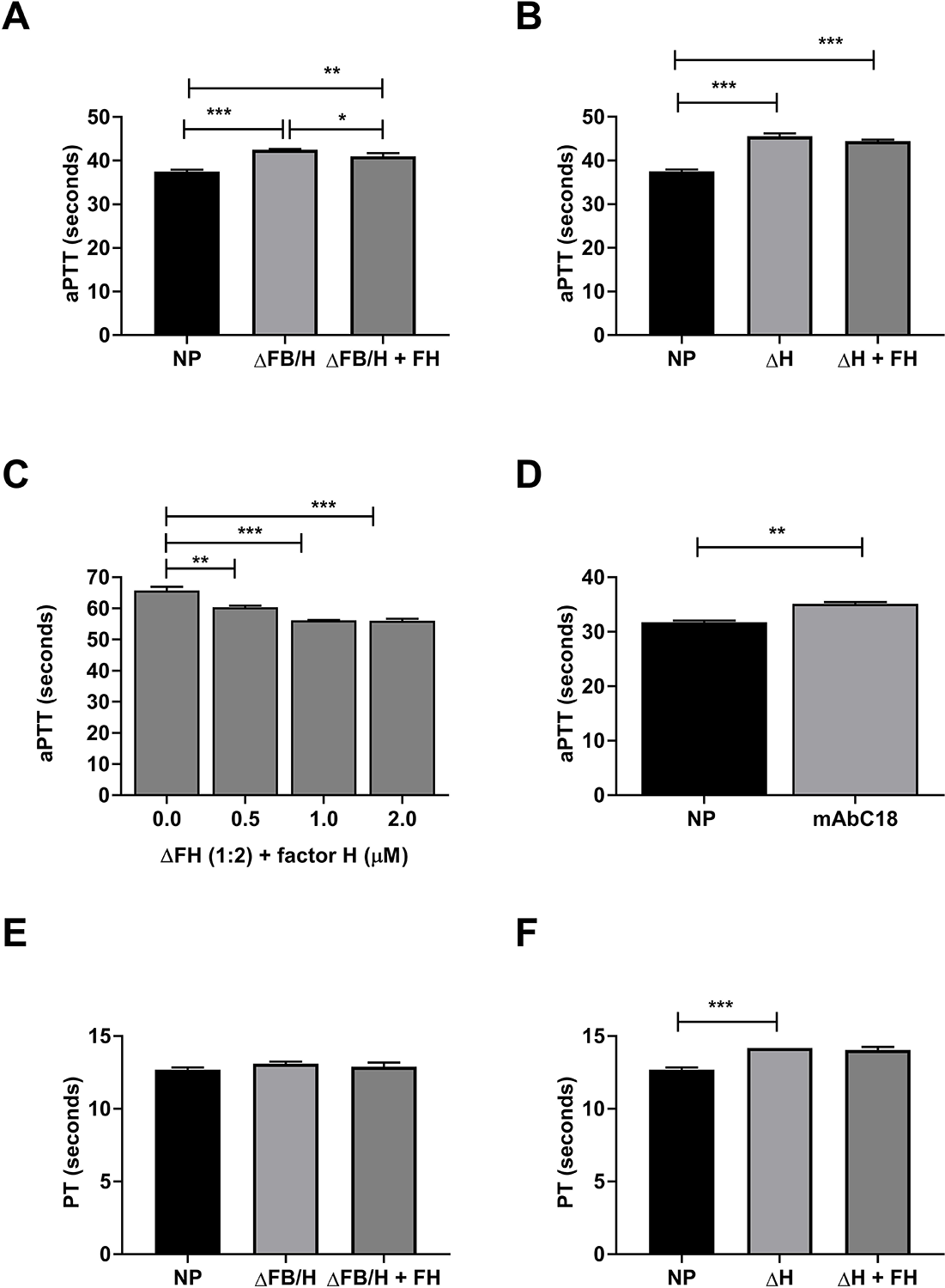
Activated partial thromboplastin time (aPTT) is elevated in human FH-depleted plasma and restored upon FH add-back. Activated partial thromboplastin time (aPTT) in FH-depleted human plasma (n=3) was elevated in **(A)** ΔFB/FH (42.53±0.15 seconds, P<0.0001) and (**B**) ΔFH (45.57±0.67 seconds, P<0.0001) in comparison to normal plasma (NP, 37.5±0.42 seconds, P=0.0002). Upon addition of 1μM FH to **(A)** ΔFB/FH aPTT was partially restored (41.03±0.7 seconds, P=0.03), but **(B)** not significantly when adding 1μM FH to ΔFH (44.43±0.35 seconds, P=0.09). (**C**) Upon ΔFH plasma dilution (1:2), aPTT decreased significantly in a concentration dependent manner when adding back FH (0.5-2μM, one-way ANOVA P<0.0001). **(D**) Addition of 1 μM mAbC18 (monoclonal anti-FH antibody recognizing its SCR20 domain) to normal plasma (NP) elevated aPTT significantly (NP 31.73±0.32 seconds vs NP+mAbC18 35.1±0.34 seconds, P=0.0022). Prothrombin time (PT) was unchanged in (**E**) ΔFB/FH (13.1±0.14seconds, P=0.25), but significantly elevated in (**F**) ΔFH (14.2±0.1seconds, p=0.004) in comparison to normal plasma (NP 12.7±0.1 seconds). Re-addition of 1μM FH did not change the prothrombin time significantly in ΔFB/FH+1μM FH (12.9±0.3 seconds, P=0.28) or to ΔFH (14.05±0.21 seconds, P=0.61). Data represented as mean±SD, n=3, and analyzed using one-way ANOVA. Representative of 3 separate experimental repeats.

The aPTT was elevated in ΔFB/FH-plasma (in the absence of complement alternative pathway) (**Figure 2A**) and ΔFH-plasma (**Figure 2B**) compared to normal human plasma (NP), confirming a prolonged clotting time in the absence of FH. Addition of FH to ΔFB/FH-plasma reduced aPTT significantly (**Figure 2A**), albeit not to NP levels, likely due to non-specific affinity-depletion or plasma dilution. Addition of FH to ΔFH-plasma did moderately reduce aPTT (**Figure 2B**); in dilute ΔFH-plasma, aPTT was reduced significantly and in a FH concentration dependent manner (**Figure 2C**).

Next, we tested whether specific blocking of FH binding domains impacts its effect on aPTT by adding an anti-FH antibody (mAbC18), which recognizes SCR20 of FH and mimics the effects of the anti-FH autoantibodies (61). Addition of mAbC18 to normal plasma resulted in elevated aPTT (**Figure 2D**). In contrast, no change in PT was observed in ΔFB/FH-plasma compared to normal human plasma (**Figure 2E**), and a moderate increase was seen in ΔFH-plasma, but FH added to ΔFH-plasma did not significantly impact PT values (**Figure 2G**).

### FH enhances rate of fibrin clot formation

The observed prolonged bleeding time and elevated aPTT in the absence of FH is compatible with dysfunctional thrombin activity and fibrin clot generation (62). Consequently, to dissect the impact of FH on fibrin clot generation, we used a pure protein assay of thrombin-mediated fibrinogen cleavage in the presence or absence of FH and monitored fibrin clot generation as turbidity change over time (**Figure 3**).

**Figure 3.**
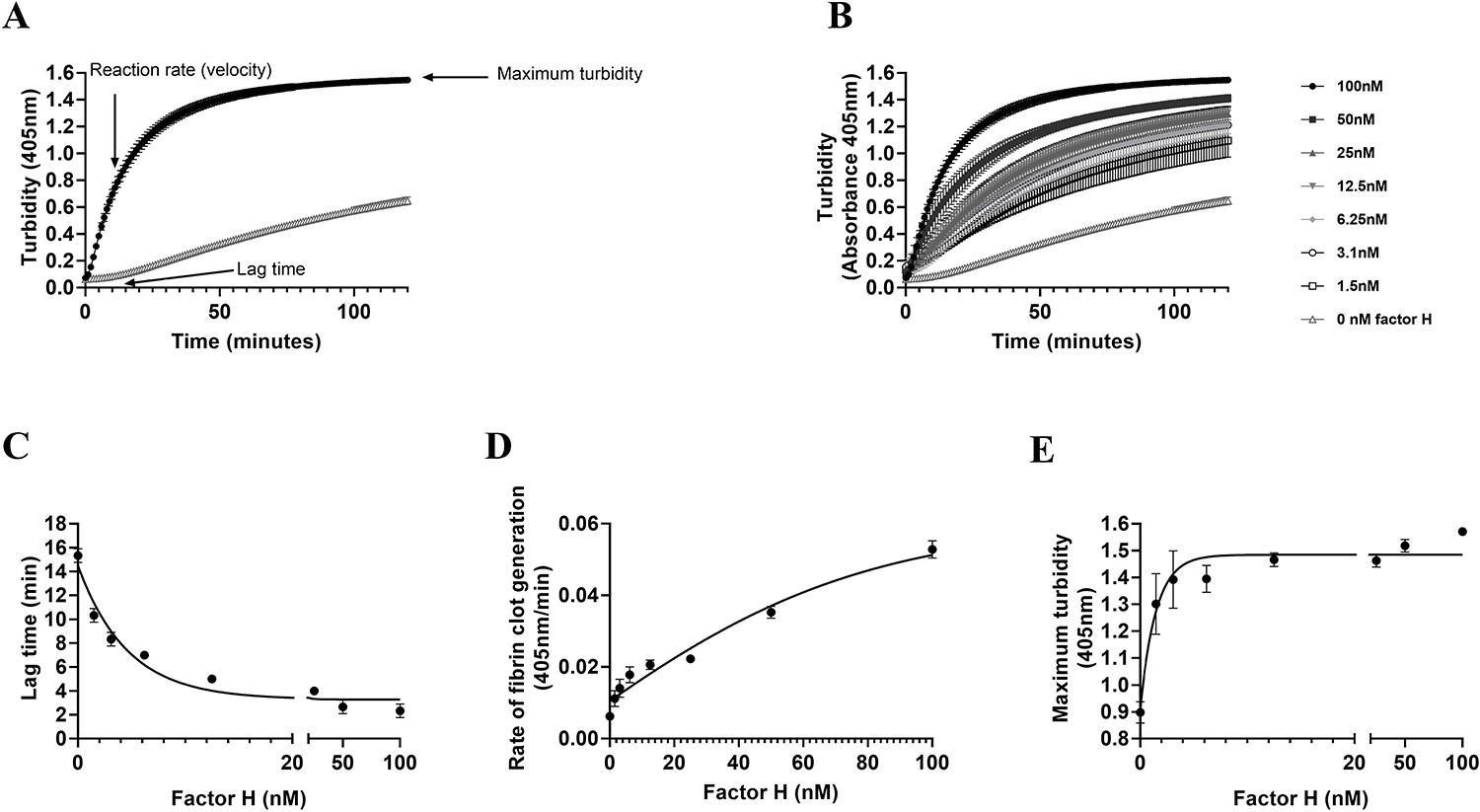
FH enhances rate of fibrin clot formation. Fibrin clot formation was monitored in a pure protein assay combining 4mg/ml (11.8μM) plasminogen-free fibrinogen with 2.5nM thrombin in 5mM Calcium buffer and FH (1.5-100nM). (**A**) Turbidity was monitored at A405nm over time to determine clotting onset (lag time, minutes), rate of fibrin clot generation (A405nm/min) and maximum turbidity (A405nm) in FH presence (100nM, •) or absence (buffer control). FH (**B**) enhanced turbidity (A405nm) over time, (**C**) shortens lag time (IC_50_: 0.64+/−0.03, P<0.0001), and (**D**) increases rate of fibrin clot generation (EC_50_: 598.1+/−100.43nM, P<0.0001) and (**E**) maximum turbidity (EC_50_: 0.8+/−0.64nM, P=0.0004) in a concentration-dependent manner. P values were analyzed using non-parametric Spearman correlation. Data represented as mean±SD, n=3. Representative of 3 separate experimental repeats.

Thrombin-mediated fibrinogen cleavage and fibrin clot formation in absence and presence of FH was monitored as turbidity (A405nm) change over time (minutes) to determine lag time, polymerization rate (velocity), and maximum turbidity, an indicator of the fibrin clot structure (**Figure 3A**). FH was added to fibrinogen and thrombin in the presence of calcium and turbidity monitored over time We detected a concentration-dependent relationship between FH and the dynamics of fibrin formation (**Figure 3B**). With increasing FH concentration, we observed a decrease in lag time (IC_50_: 0.64+/−0.03, **Figure 3C**), increased rate of fibrin clot generation (velocity) (EC_50_: 598.1+/−100.43nM, **Figure 3D**) and maximum turbidity (EC_50_: 0.8+/−0.64nM, **Figure 3E**). The results suggest that FH affected thrombin-mediated cleavage of fibrinogen into fibrin and protofibril formation, the rate of the reaction and fiber formation, and the structure of the fibrin clot.

### FH alters fibrin clot structure and fiber density

Thicker fibers are associated with more opaque gels and increased maximum turbidity (63–65). Having shown that FH affected fibrin clot maximum turbidity, we next wanted to establish whether FH affects fibrin clot structure. To explore this, we generated pure fibrin clots by incubating thrombin with fibrinogen in the presence or absence of FH and imaged the matured fibrin clot by fluorescence microscopy (**Figure 4**).

**Figure 4.**
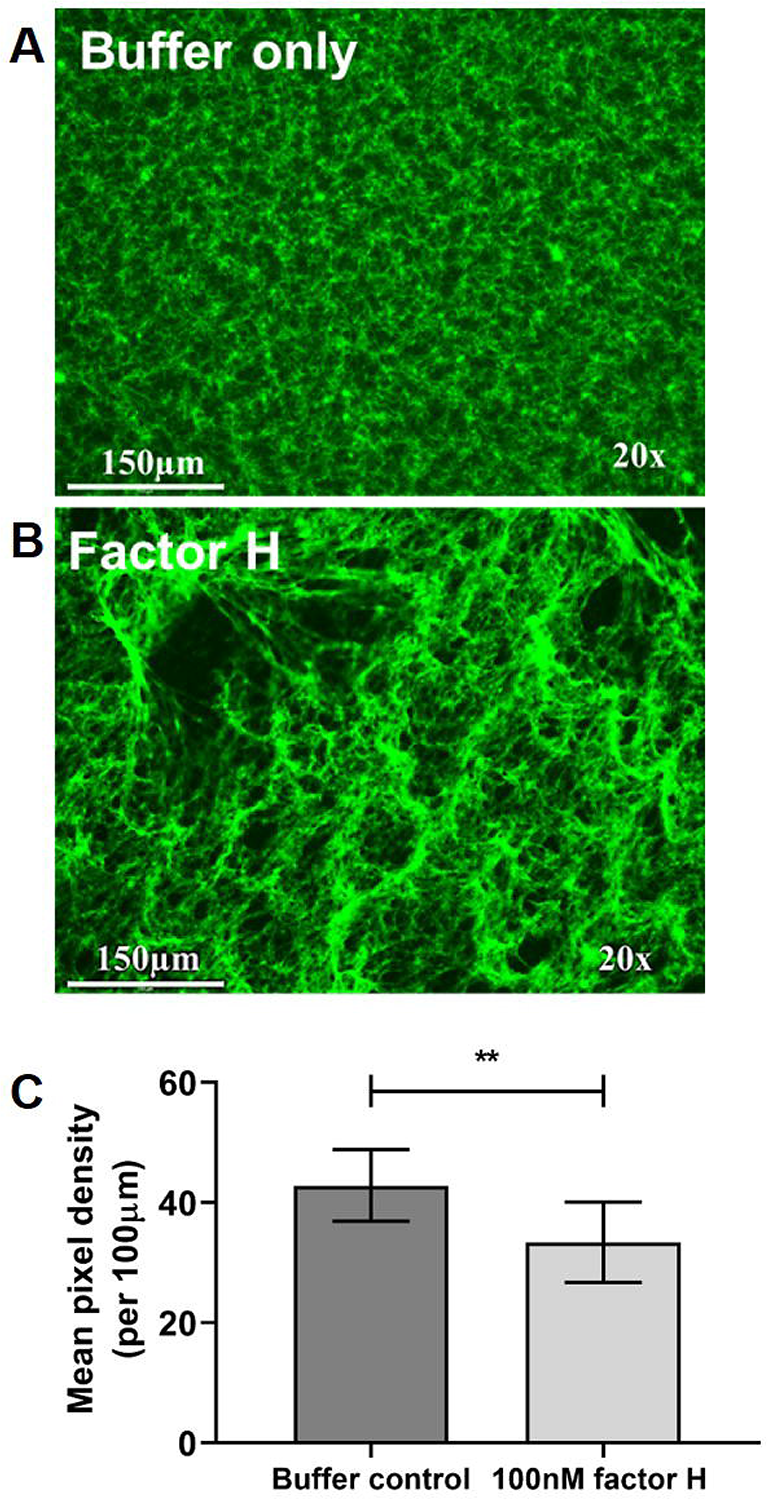
FH alters fibrin clot structure and fiber density. A fibrin clot was generated by combining 11.8μM plasminogen-free fibrinogen containing 10%(w/v) fibrinogen-AF488 with 2.5nM thrombin in 5mM Calcium buffer in the presence (100nM) or absence of FH, and left to mature for 2 hours. The fibrin clot was visualized by fluorescence microscopy (EVOS™ M7000 Imaging System) at 20X magnification. Images (scale bar 150μm) depict the fibrin clot structure in the (**A**) absence (buffer control) or (**B**) presence of 100nM FH. (**C**) Fiber density was determined by ImageJ Version 1.43 using the plug-in grid. Mean pixel density was calculated by counting the number of fibers crossing 25 pre-defined 100μm lines placed in each image from a total of 12 images taken in different areas of the clot. Mean pixel density is decreased in the presence of FH (33±7 mean pixel density, P=0.0013) compared to buffer control (43±6 mean pixel density). P value determined by non-parametric Mann– Whitney test. Data represented as mean ±SD. Image and data representative of 9 experimental repeats.

The presence of FH altered the fibrin clot structure compared to the buffer control (**Figure 4A**). In the presence of FH, clots comprised large fibers separated by more numerous and larger pores (**Figure 4B**) and fiber density was significantly reduced (**Figure 4C)**, suggesting a role of FH in the formation and structure of the fibrin gel (13,66).

### FH enhances thrombin-TM and thrombin-mediated PC activation

Given the versatile roles of thrombin in hemostasis, we next wanted to determine whether FH also affected thrombin-mediated PC activation in the presence or absence of TM (67–69). Thrombin binding to its cofactor TM increases APC generation several-fold (70,71) and we have previously shown that FH also binds TM with high affinity (25). To test whether FH affects PC activation, we tested interactions between PC, thrombin, TM and FH (**Figure 5**). We observed a FH-dependent increase in PC activation and determined the affinity constant Kd for thrombin-TM complex (•Kd 1.6±0.2nM FH, P <0.0001) and thrombin-only (◻, Kd 29.1±2.8nM FH, P=0.0003) mediated PC activation (**Figure 5A**). FH induced significantly faster PC activation kinetics when titrating PC in the presence or absence of FH with TM-thrombin (• 100nM FH, KM: 80±28nM PC compared to ◻ buffer control, KM: 187±105nM PC; P=0.004) (**Figure 5B**) or thrombin-only (• 100nM FH, KM: 199±38nM PC; or ◻ buffer control, KM: ~752±99nM PC, ~approximation as the reaction doesn’t reach saturation) (**Figure 5C)**. Next, we wanted to determine whether the enhancing effect of FH can be blocked using a mAb targeting the FH SCR20 region. The mAb C18 caused a partial reduction in FH-mediated increased PC activation rate (Buffer control: 0.059±0.002 vs 100nM FH: 0.128±0.001 nM/min vs 100nM FH + mAbC18: 0.121±0.001 nM/min, P<0.01; **Figure 5D**); interactions of FH with thrombin-TM (**Figure 5E**) and thrombin-TM456 were not reduced significantly in the presence of mAbC18 (Buffer control: 0.038±0.0004 vs 100nM FH: 0.081±0.003 nM/min vs 100nM FH + mAbC18: 0.075±0.001 nM/min, P=0.058). While FH enhanced the rate of PC generation by (**Figure 5F**) thrombin alone, the anti-FH antibody showed no effect (Buffer control (−) 0.02±0.0007 vs 100nM FH (FH) 0.039±0.0005 nM/min vs 100nM FH (FH) + mAbC18 0.039±0.001 nM/min, P<0.95).

**Figure 5.**
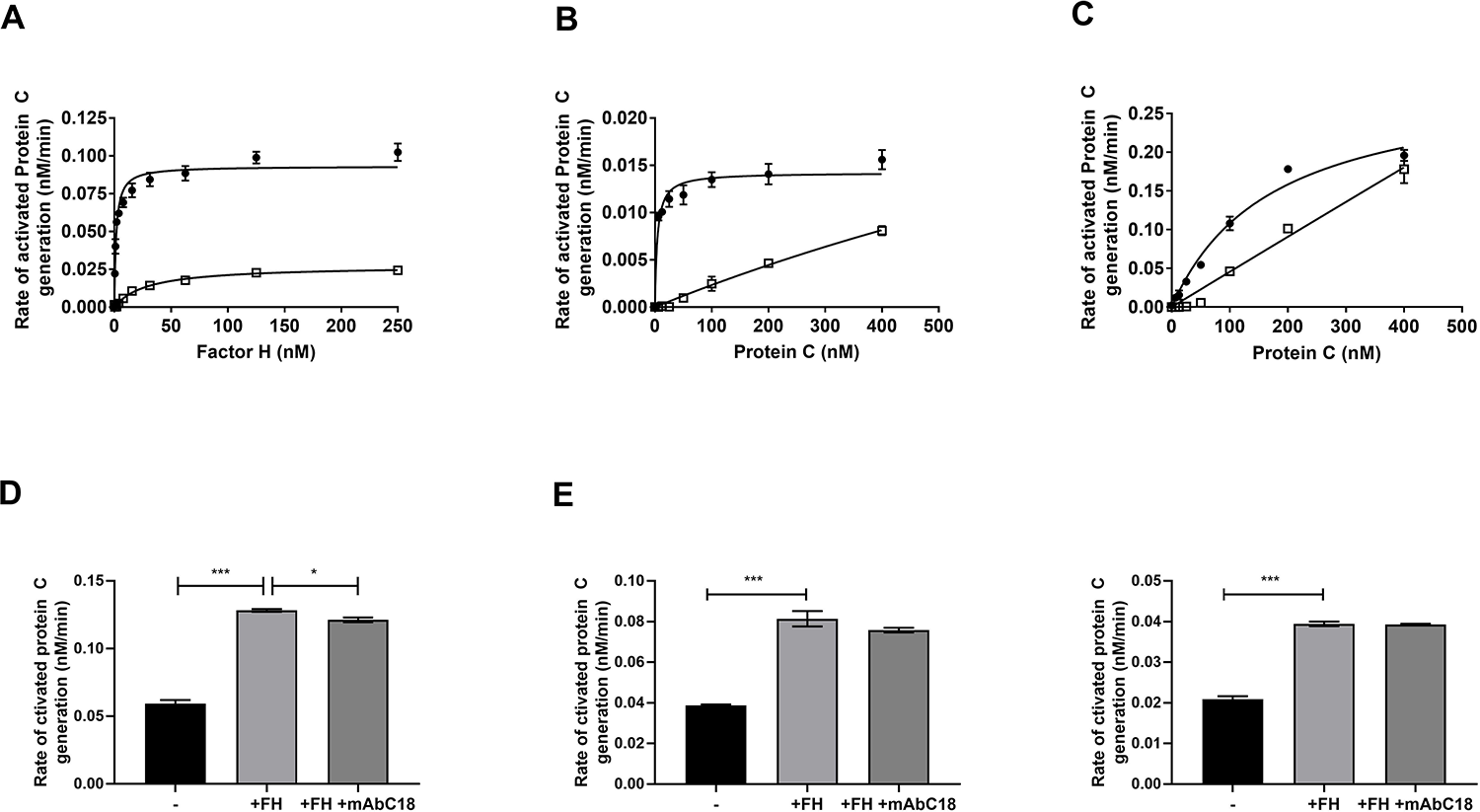
FH enhances PC activation. FH was added to 1.75nM thrombin and 100nM PC, in the presence or absence of 10nM TM in 3mM Calcium buffer. The reaction was stopped after 2 hours by adding 1U/well Hirudin. APC generation was quantified by monitoring chromogenic APC substrate cleavage (A405nm) over time (**A**). FH (0.5-125nM) enhanced PC activation determined as binding constant Kd for thrombin-TM (•Kd 1.6±0.2nM FH, P <0.0001) or thrombin-only (◻, Kd 29.1±2.8nM FH, P=0.0003) in a FH concentration-dependent manner. (**B**) FH (100nM) induced significantly faster PC activation kinetics when titrating PC with added thrombin-only (• 100nM FH: KM 199±38nM PC; or ◻ buffer control: KM ~752±99nM PC, ~approximation as the reaction doesn’t reach saturation) or by **(C**) thrombin-TM (• 100nM FH: KM 80±28nM PC compared to ◻ buffer control: KM 187±105nM PC; P=0.004). P values were analyzed by Spearman correlation. FH (100nM, light grey) significantly enhances the rate of PC generation by (**D**) thrombin-TM (black, P<0.0001), while a FH SCR20 blocking antibody (100 nM mAbC18) partially reduces the FH effect (dark grey; buffer control (−) 0.059±0.002 vs 100nM FH 0.128±0.001 nM/min vs 100nM FH + mAbC18 0.121±0.001 nM/min, P<0.01), and also enhances PC generation mediated by (**E**) thrombin with recombinant TM456 fragment (500nM, black, P<0.0001), while a FH SCR20 blocking antibody (100 nM mAbC18) did not significantly reduce the FH effect (dark grey; buffer control (−) 0.038±0.0004 vs 100nM FH 0.081±0.003 nM/min vs 100nM FH (FH) + mAbC18 0.075±0.001 nM/min, P=0.058). FH also enhanced (**F**) thrombin-only (black, P<0.0001) mediated PC generation, but the FH SCR20 blocking antibody (100 nM mAbC18) did not significantly reduce the FH effect (dark grey; buffer control (−) 0.02±0.0007 vs 100nM FH (FH) 0.039±0.0005 nM/min vs 100nM FH + mAbC18 0.039±0.001 nM/min, P=0.95). Data represented as mean±SD, n=3. P values were analyzed by Mann Whitney test. Representative of 3 experimental repeats.

### FH binding interaction with thrombin and affinity analysis

Next, we wanted to establish whether the FH modulation of coagulation and anticoagulation was a consequence of direct binding to thrombin and fibrinogen in addition to its known interaction with TM (22). To achieve this, we used ELISA to investigate complex formation between FH, thrombin or prothrombin and determined the binding affinity of FH with thrombin or fibrinogen by SPR analysis (**Figure 6**).

**Figure 6.**
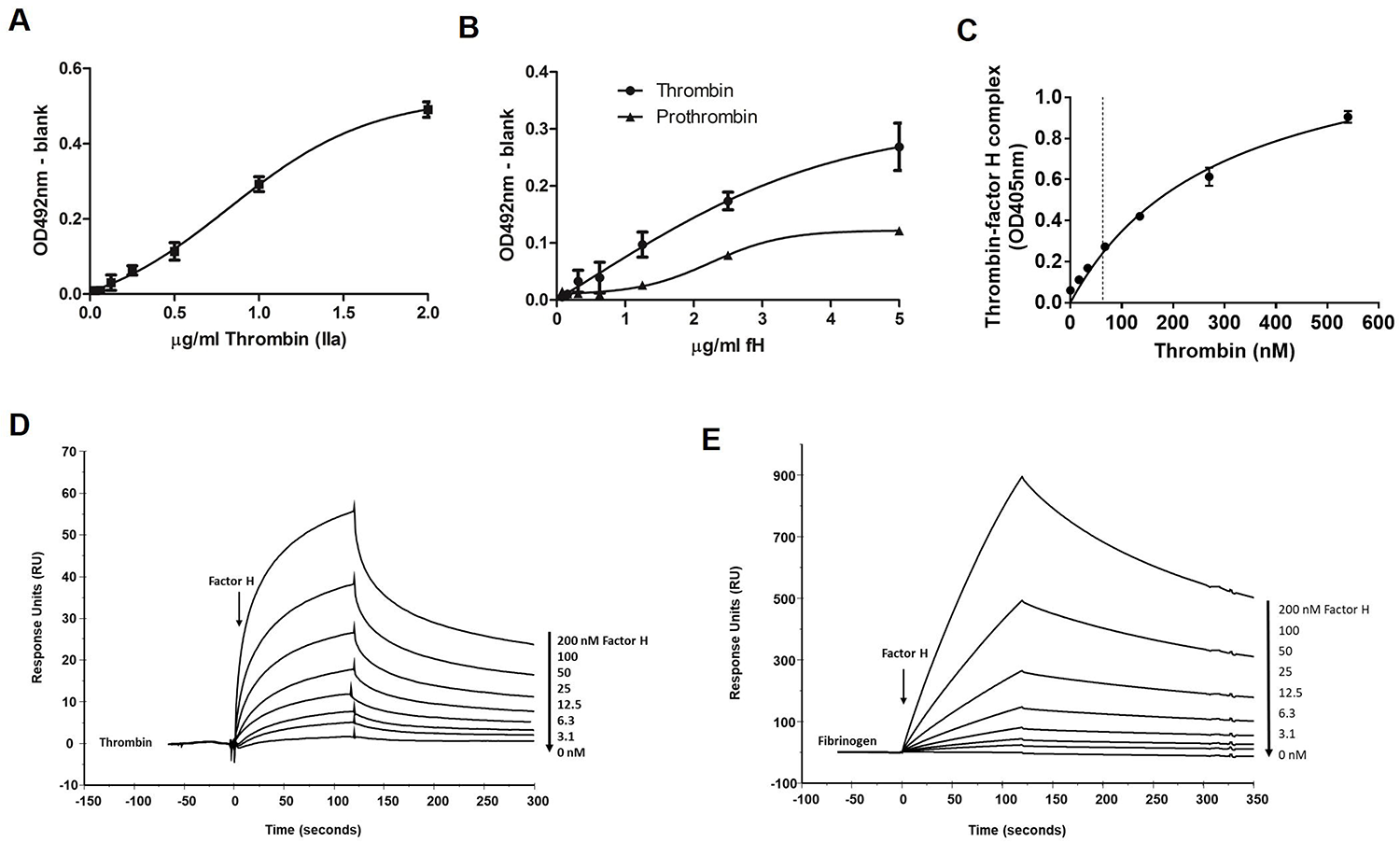
FH binds thrombin with nanomolar affinity and forms a measurable FH-thrombin complex in plasma. (**A**) Thrombin (0.06-2 g/ml) binding to microtitre plate adsorbed FH (◾) was measured using a polyclonal anti-thrombin antibody. (**B**) FH (0.15-5 g/ml) binding to microtitre plate adsorbed PPACK-thrombin (•) and to a lesser extend to prothrombin (▴) was measured using a polyclonal anti-FH antibody. (**C**) Normal EDTA-plasma (NP) was spiked with PPACK-thrombin (1 M) and FH-thrombin complex (dashed line) was quantified in ELISA at 62nM FH-thrombin, using antibody-captured FH and a thrombin standard curve (17-540nM). Binding affinity of FH (3.1-200 nM) to immobilized (**D**) thrombin (1000U) was determined at KD: 29.7±1.08nM and for (**E**) fibrinogen at KD 38.2±0.93 nM. Affinity was calculated using a 1:1 binding Langmuir model and Biacore T200 software v3.1. Data represented as mean ±SD, n=3.

To elucidate a direct interaction between thrombin and FH, we either incubated thrombin with FH captured on microtiter plates (**Figure 6A)** or FH with thrombin or prothrombin captured on a microtiter plate (**Figure 6B**), and found a dose-dependent interaction between FH and thrombin in both orientations (**Figure 6A and 6B**). Prothrombin did not bind FH with appreciable affinity (**Figure 6B**), which was also confirmed by SPR analysis (data not shown). The in vitro complex formation between FH and thrombin in plasma was investigated by ELISA by adding thrombin at supraphysiological concentration to normal plasma and measuring FH-thrombin complex in the lower nanomolar range (**Figure 6C**). SPR analysis was used to further investigate the interaction and to determine the binding strength of FH with thrombin. PPACK-thrombin was immobilized onto the chip surface and varying FH concentrations were flown over the surface. FH binding affinity with immobilized thrombin was KD 29.7±1.08nM (**Figure 6D**). Furthermore, we determined that FH binding affinity with immobilized fibrinogen was KD 38.2±0.93 (**Figure 6E**).

Finally, we tested whether thrombin affects FH and its regulatory activity in the complement system (**Figure 7**). We showed that thrombin did not proteolytically cleave FH (**Figure 7A**); we observed only minor FH degradation at the highest thrombin concentrations (>250nM) after 90-minute incubation (**Figure 7B**). In normal human serum, thrombin did not itself have any significant regulatory effect on complement mediated hemolysis (**Figure 7C**), and did not show any effect on FH-mediated regulation of hemolysis at physiological concentration (**Figure 7D**). While we observed some moderate effect of supraphysiological thrombin on FH regulation (**Figure 7E**), this was also seen with catalytically inactive PPACK-thrombin (**Figure 7F**), and likely due to competitive binding of thrombin with FH and its ligands. We confirmed by SDS-PAGE and mass spectrometry analysis that FH exposed to supraphysiological thrombin was not degraded and maintained its intact molecular weight (**Supplementary Figure 2).**

**Figure 7.**
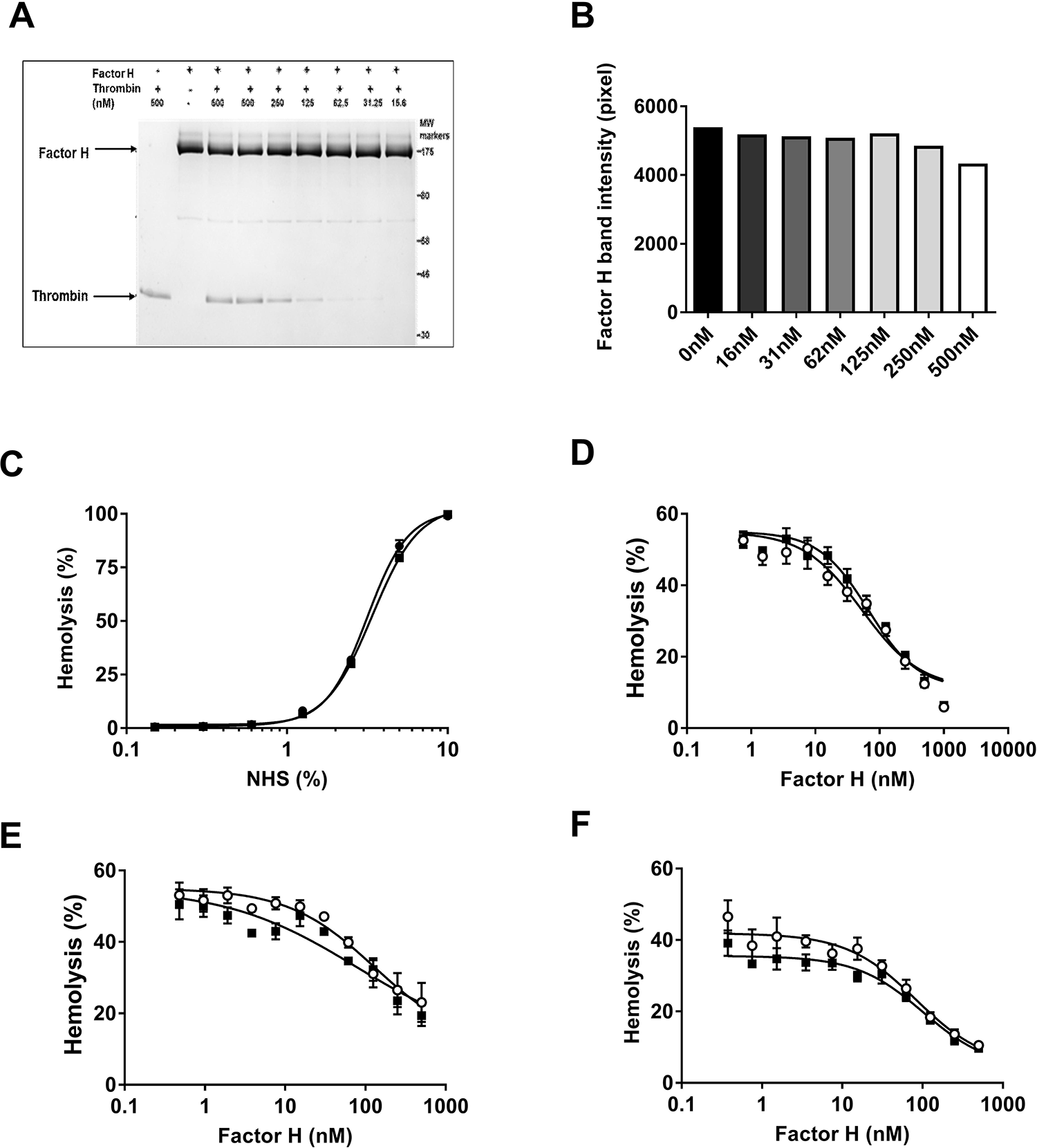
Thrombin does not cleave or affect FH complement regulation. **(A)** Thrombin (15-500nM) was incubated with FH (2ug per lane) and incubated for 90 minutes and FH band resolved on a 4-12% gradient gel and visualized with Coomassie staining. **(B)** Thrombin does not proteolytically cleave FH. Minor FH band degradation is observed at very high thrombin concentrations above 250nM after 90-minutes incubation. **(C)** Thrombin (500nM) did not have any regulatory effect on complement mediated hemolysis in normal human serum (NHS) (EC_50Thrombin_: 3.39±0.11 %NHS vs EC_50Control_: 3.15±0.11 %NHS, P=0.1) and **(D)** did not show any effect on FH-mediated regulation of hemolysis at physiological thrombin (10nM) concentration (EC_50Thrombin (500nM)_: 190.7±92.29 vs EC_50Control_: 206.2±58.9nM FH, P=0.99). Supraphysiological concentration of **(E)** thrombin (500nM) moderately decreased FH regulation (EC_50Thrombin (500nM)_: 115.2±25.73 vs EC_50Control_: 78.73±20.97nM FH, P=0.99) and **(F)** PPACK-thrombin (500nM) concentration (EC_50Thrombin-PPACK (500nM)_: 120.7±37.4 %NHS vs EC_50Control_: 85.75±24.64 %NHS, P=0.1).

We concluded that FH is a thrombin ligand and mediates its pro- and anticoagulant activities, and that thrombin does not significantly affect FH complement regulatory ability.

## Discussion

FH is a key regulator of the complement alternative pathway (72), downregulating complement amplification by acting as a cofactor for FI-mediated cleavage and inactivation of opsonin C3b. FH deficiency or dysfunction is characterized by excessive complement activation, resulting in pathology also associated with coagulation dysfunction, for instance TMA in aHUS.

FH deficiency in *CFH^−/−^* mice leads to excessive complement activation and spontaneous development of glomerulonephritis (34). We hypothesized that FH may also influence the clotting ability of *CFH^−/−^* mice and showed that absence of FH resulted in prolonged tail bleeding in vivo. We observed evidence of stop-start bleeding in *CFH^−/−^* mice with prolonged bleeding, suggestive of a reduction in fibrin clot formation or its stability. Plasma levels of soluble TM were marginally increased in *CFH^−/−^* mice compared to WT levels, indicating endothelial damage (18), and plasma-Hb, a marker of intravascular hemolysis (73,74), was also slightly increased in *CFH^−/−^* mice. However, plasma levels of thrombin-antithrombin (TAT) complex, a marker of *in vivo* thrombin generation was not increased in *CFH^−/−^* mice, suggesting no pathological activation of the coagulation pathway and that prolonged tail bleeding was not due to consumption of coagulation factors as may be seen for example in disseminated intravascular coagulation (DIC) but not TMA (75).

As there was no evidence of increased activation of coagulation we wanted to determine whether plasma FH deficiency was associated with a defect in coagulation pathways. We therefore measured activated partial thromboplastin time (aPTT) and prothrombin time (PT) in both FH-depleted or FH and FB double-depleted human plasma, representing the presence or absence of the alternative complement pathway capability, respectively. FH-depleted plasma showed a prolonged aPTT compared to similarly treated normal plasma; this was partially restored upon re-addition of physiological concentrations of FH indicating that FH directly contributed to the generation of a fibrin clot. FH depletion modestly elevated the PT compared to normal plasma, however no significant reduction was observed upon re-addition of FH, possibly due to elevated thrombin generation due to tissue factor activation, overwhelming FH effects.

Elevated aPTT and PT is likely caused by a process that influences elements of the common pathway (76). FH is a cofactor for complement serine protease FI and because there is considerable structural homology between the FI and thrombin (30), we investigated the possible effects of FH on thrombin-mediated coagulation functions. We hypothesized that the mechanism by which aPTT was prolonged may be directly linked to thrombin-mediated fibrin clot formation. Therefore, we assessed the role of FH in fibrin clot generation and clot structure in pure protein assays. We identified a direct role for FH in coagulation by enhancing thrombin-mediated fibrin formation a dose-dependent manner at low nanomolar concentrations. Increasing FH concentrations both increased the polymerization rate and decreased lag time in clot turbidity assays, suggesting an accelerated association of fibrin oligomers into protofibrils and then into fibers (77), possibly affecting lateral aggregation (12,78). Increased lateral aggregation leads to larger fibers and less branching points with larger pores (77–81), which we observed when imaging a pure fibrin clot formed in the presence of FH. We demonstrated decreased fiber density in the presence of FH, which would allow for greater clot permeability and potential access of fibrinolytic components (82). Further study is needed to dissect the impact of FH on factor XIIIa mediated stabilization of fibrin, the susceptibility to fibrinolysis (77,78), fibrin clot diffusivity (13,66), or access of fibrinolytic plasmin (13,83).

To confirm whether the observed effect on fibrin clot generation and structure is a result of FH binding interaction with fibrinogen and/or thrombin we performed binding interaction analysis and found that FH binds with strong affinity with fibrinogen and thrombin with affinity in the low nanomolar range. FH is an abundant plasma protein with fluid phase concentrations in the micromolar range (0.7-3.6 μM) (84,85). The nanomolar affinity constant suggest that plasma FH could readily bind thrombin on its formation and also to circulating fibrinogen. The binding affinities are also in contrast to the weaker affinity of FH with C3b, which is in the micromolar range (86). Additionally, we show that endogenous plasma FH forms stable FH-thrombin complexes in the nanomolar range when excess thrombin is added to EDTA-plasma.

A key aspect of thrombin function is its ability as a regulator of its own formation, thrombin acts as an anticoagulant via the thrombin-TM mediated activation of protein C (70,71). We have previously demonstrated that FH can bind to TM with nanomolar affinity (22), and our data showing nanomolar affinity for FH-thrombin binding, which led us to investigate FH in thrombin-TM mediated PC activation. We observed a FH concentration dependent increase in the rate of PC activation, decreasing KM, by both the thrombin-TM complex, and to a lesser extent, by thrombin alone. Thrombin alone is not considered a physiological activator of PC (69,87–89), but we observed a several-fold enhanced thrombin-mediated activation of PC in the presence of FH. The physiological significance of FH enhacing thrombin-only mediated PC activation needs further investigation, but may be sufficient to support limited activation of PC at TM-restricted sites, such as the fibrin clot microenvironment and thus support localized clot formation.

Overall, FH binding to TM (22) and thrombin, its impact on PC generation as well as thrombin-mediated fibrin clot generation supports the hypothesis that FH acts as a direct cofactor for thrombin. The presence of FH enhanced the rate of thrombin-catalyzed fibrin formation and increased the catalytic efficiency of the thrombin-TM complex in the activation of PC. While thrombin interaction with its substrates is normally mediated by its active site and its exosites (7,90), it is known that cofactors regulate thrombin activity through competition for exosite binding, allostery (6) and cell surface localization (4,6,62). Interestingly, FH, while present in plasma, is both a fluid phase and cell surface regulator. Surface binding of FH, for instance, to the endothelium (91), is integral to complement regulation; binding occurs through interactions with glycosaminoglycans (72), phospholipids (92,93) and opsonin C3b (94,95). Furthermore, we showed that antibody-blocking FH SCR20 domain which mimics the effects of the anti-FH autoantibodies in disease (61) affects its thrombin modulating function in both procoagulant (aPTT) and anticoagulant (APC generation) assays, indicating that the SCR20 domain is important for FH binding with thrombin and thrombin-TM. FH SCR20 is relevant to FH surface binding and regulation of complement amplification; FH could promote surface localization of thrombin and by binding with fibrinogen (49,96), clot components, and its localization in the fibrin clot (97–99). Our study showed that FH is a high affinity ligand for thrombin, but not a proteolytic substrate, suggesting a role as a cofactor modulating thrombin activity in both its procoagulant role in the cleavage of fibrinogen, and its anticoagulant role with TM in the activation of PC. Further study is required to determine whether this is due to distinct allosteric forms of thrombin and FH-thrombin complex, similar to the slow to fast form transition of thrombin with Na+ binding, regulating its anticoagulant and procoagulant activities (100,101).

Finally, while thrombin supports complement activation in vitro (43,44), we found no significant impact of thrombin on FH regulatory role in the complement system. Our study highlights the importance of molecular interactions between complement, coagulation and anticoagulation pathways and the potential impact of their dysregulation in diseases where complement and coagulation dysfunction or deficiency contributes to pathology.

## Acknowledgements

We would like to thank Dr K. Allen-Redpath for help with establishment of the tail bleeding assay, and Ms R. Burke (MPharm) for help with the development of the thrombin-FH binding assay. This work was funded by internal research funding from the School of Pharmacy and Pharmaceutical Sciences, Cardiff University (2017-2021, MH) and ISSF-Welcome Trust Seeds for seed award 516642 (2018-2021, MH) and NISCHR award HF-11-22 (2012-2016, MH). PVJ is supported by Health and Research Wales CRTA 16-05.

## Authorship Contributions

Conceptualization: M.H.; methodology and experimental design, M.H., G.M.; experiments and data analysis, M.H., G.M., G.D., R.V., animal studies, T.R.H.; formal analysis and consolidation, M.H. and G.M.; writing—original draft preparation, M.H. and G.M., writing—interpretation, review and editing, M.H., G.M., R.P., B.P.M, P.W.C, P.V.J, supervision, M.H.; project administration, M.H., All authors have read and agreed to the published version of the manuscript.

Correspondence: Meike Heurich, School of Pharmacy and Pharmaceutical Sciences, Cardiff University, United Kingdom;

