## Supplementary material for "Complement Regulator Factor H is a Cofactor for Thrombin in both Pro- and Anticoagulant Roles"

**Materials**

RecombiPlastin 2G (RTF 2G, or PT reagent), Calcium Chloride 0.020M (CaCl_2_) and SnythASil (APTT reagent) were purchased from HemosIL, for use on the MC10 coagulometer only. Human alpha-thrombin, catalytically inactive PPACK-thrombin, human protein C, human activated protein C and plasminogen-depleted fibrinogen (FXIII is 0.08IU/ml 2.9 mg/ml, Clauss Fibrinogen) were all purchased from Enzyme Research Laboratories (Swansea, UK). Fibrinogen Alexa Fluor 488 was purchased from Invitrogen. Human soluble recombinant thrombomodulin (amino acid Ala19-Ser515 (Ala473Val), with a C-terminal 6-His tag, apparent molecular weight 80-105 kDa, determined by SDS-PAGE under reducing conditions) was purchased from R&D Systems (Biotechne, UK). Recombinant TM456 was a kind gift from Prof J. Huntington, Cambridge University, UK). Activated protein C substrate was from Biophen CS21(66), Quadratech Ltd. Sheep erythrocytes were purchased from TC Biosciences, and normal human serum was collected from healthy donors and pooled. Monoclonal anti-FH antibody clone C18/3, which recognizes SCR20 of fH was purchased from Enzo Life Sciences (UK). Mouse anti-factor H monoclonal antibody clone 35H9, mouse anti-factor H monoclonal clone 10-15 and rabbit anti-factor H polyclonal antibody were all made in-house. Donkey anti-mouse IgM antibody and anti-sheep IgG polyclonal antibodies were from Jackson ImmunoResearch Inc (USA). Hirudin (1unit= 1μl) and HRP substrate (OPD, O-phenylenediamine dihydrochloride) were from Sigma-Aldrich Company Ltd (UK). Non-fat dried milk was from The Co-Operative Group Ltd. Polystyrene plates (MaxiSorpTM) were from Fisher Scientific UK Ltd. 96 well plates flat bottomed and U-bottomed were purchased from Fisher Scientific UK Ltd, precast 4-12% Tris-glycine gradient gels were from Life Technologies Ltd. Glass slides and coverslips were purchased from Dixon Science (UK). Biacore T200 and series S Sensor Chip CM5 was from GE Healthcare.

**Supplementary methods**

**Animal studies**

All animals were housed in conventional cages at the SPF facility, Cardiff University, Cardiff, United Kingdom. Where needed animals were anaesthetised using *Isoflurane.* All animal studies were carried out under the authority of project licence (PPL) 30/3365 granted by the UK Home Office. Both male and female animals were used in our studies, data presented here are male only. Wild-type (WT) mice (C57/Black6) were purchased from Envigo, mice with factor H deficiency (CFH^−/−^) on a C57/Black6 background have been described previously (1) were a kind gift of Prof. Matthew Pickering were bred and maintained in our facility.

**Tail bleeding time**

Mice were weighed and anaesthetised using isoflurane. Animals were placed in prone position. A 3-mm portion of distal tail tip was amputated with a scalpel. The tail was immediately immersed in a 50-mL Falcon tube containing isotonic phosphate buffered saline pre-warmed in a water bath to 37 °C.

The position of the tail was vertical with the tip positioned about 2 cm below the level of the body. Each animal was monitored for 20 min even if bleeding ceased, in order to detect any re-bleeding. Bleeding time was determined using a stop clock. If bleeding on/off cycles occurred, the sum of bleeding times within the 20-min period was used. The experiment was terminated at the end of 20 min to avoid lethality during the experiment as required by the authorised licenced protocol.

**Mouse plasma analysis**

At the end of experiment, animals were killed by exsanguination via cardiac puncture under surgical plane anaesthesia, death was confirmed by cervical dislocation. Whole blood was drawn into citrate solution and processed as follows.

Whole blood underwent centrifuging at 4500 r/min for 5 minutes at room temperature. Plasma supernatant was removed and stored at -80°C until further analysis. Mouse plasma hemoglobin were measured spectrophotometrically by absorbance at 405nm using a (FLUOstar® Omega, BMG Labtech).

**Plasma protein purification and affinity depletion of plasma**

FH protein was eluted from immunoaffinity columns with 0.1M Glycine/HCl pH 2.5 and dialysed into 10mM HEPES, 150mM NaCl, pH7.4 and stored in aliquots at -80 °C. The factor used in Biacore studies was gel filtered into assay buffer using a Superdex 200 10/300 column (GE Healthcare) immediately prior to analysis. To generate depleted plasma, blood samples from at least three healthy volunteers were anticoagulated with sodium citrate, plasma supernatant collected after centrifugation at 4000rpm for 15 minutes (twice), pooled and stored at −80°C. Citrated plasma was factor H-depleted by flowing over sequentially anti-factor B (clone JCI, in house) and anti-factor H (clone 35H9, in house) immobilised to Sepharose columns (Hi-Trap HP-NHS, GE Healthcare). Factor B depletion is required to prevent alternative pathway mediated complement consumption in the absence of factor H. The factor H depleted plasma termed ΔFH plasma, was collected aliquots were stored at −80°C.

**Determination of FH and coagulation factor levels in FH-depleted plasmas**

To confirm depletion of FH, we used HycultBiotech (NL) HK342-02 ELISA kit to quantify FH in citrated plasma. In normal plasma (NP), FH was within its normal range at 246±4 g/ml, and below ELISA kit detection levels in FH-depleted plasmas (ΔBH and ΔH).

**Determination of factor H proteolysis by active thrombin**

FH (2μg/ml) is incubated the presence or absence of thrombin (500nM) for 1 hour at 37◦C, the reaction stopped with non-reduced or DTT-reduced SDS buffer and bands visualised on SDS-PAGE by Coomassie stain (4-12% Tris-Glycine gradient gel).

Factor thrombin was excised and subjected to in-gel analysis of factor H in the presence of thrombin was subjected to MALDI analysis via external service (sample +/- desalt C4zt, elute 50% MeCN/0.1%TFA. DHAP matrix. 20-150k m/z, Bruker Ufx LP10-50) to identify potential degradation of factor H in the presence of thrombin. Maldi-ISD analysis was used to interrogate N- terminal region.

**Hemolysis assay with active thrombin added to normal human serum**

Alpha-thrombin (500nM) is added to a dilution of 10% normal human serum (NHS). NHS serial dilutions in complement fixing buffer (CFD, Oxoid, UK) is added to a 2% suspension of Amboceptor-sensitised sheep erythrocytes (ShEA) and incubated at 37 °C for one hour followed by centrifugation at 1500rpm for 5 minutes. Hemoglobin content in supernatant is determined by measuring absorbance at A405nm.

**Hemolysis assay of factor H regulating complement in normal human serum**

Factor H (0.49-500nM) in the presence or absence of 0.625% normal human serum (NHS) with added PPACK-thrombin or alpha-thrombin (500nM), yielding 40-50% lysis and a 2% suspension of Amboceptor-sensitised sheep erythrocytes (ShEA) in complement fixing buffer. Cell suspension is incubated at 37 °C for one hour followed by centrifugation at 1500rpm for 5 minutes. Hemoglobin content in supernatant is determined by measuring absorbance at A405nm.

**Calculation of percentage Hemolysis**

Percentage hemolysis is calculated using a negative control (NC): 2%ShEA with added CFD buffer (0% lysis), and positive control (PC): %ShEA with added 0.01% Tween in H_2_O (100% lysis) as follows:

% Hemolysis = 100*(A405nm sample-A405nm NC) / (A405nm PC - A405nm NC).

**Supplementary results**

**Supplementary Figure 1**

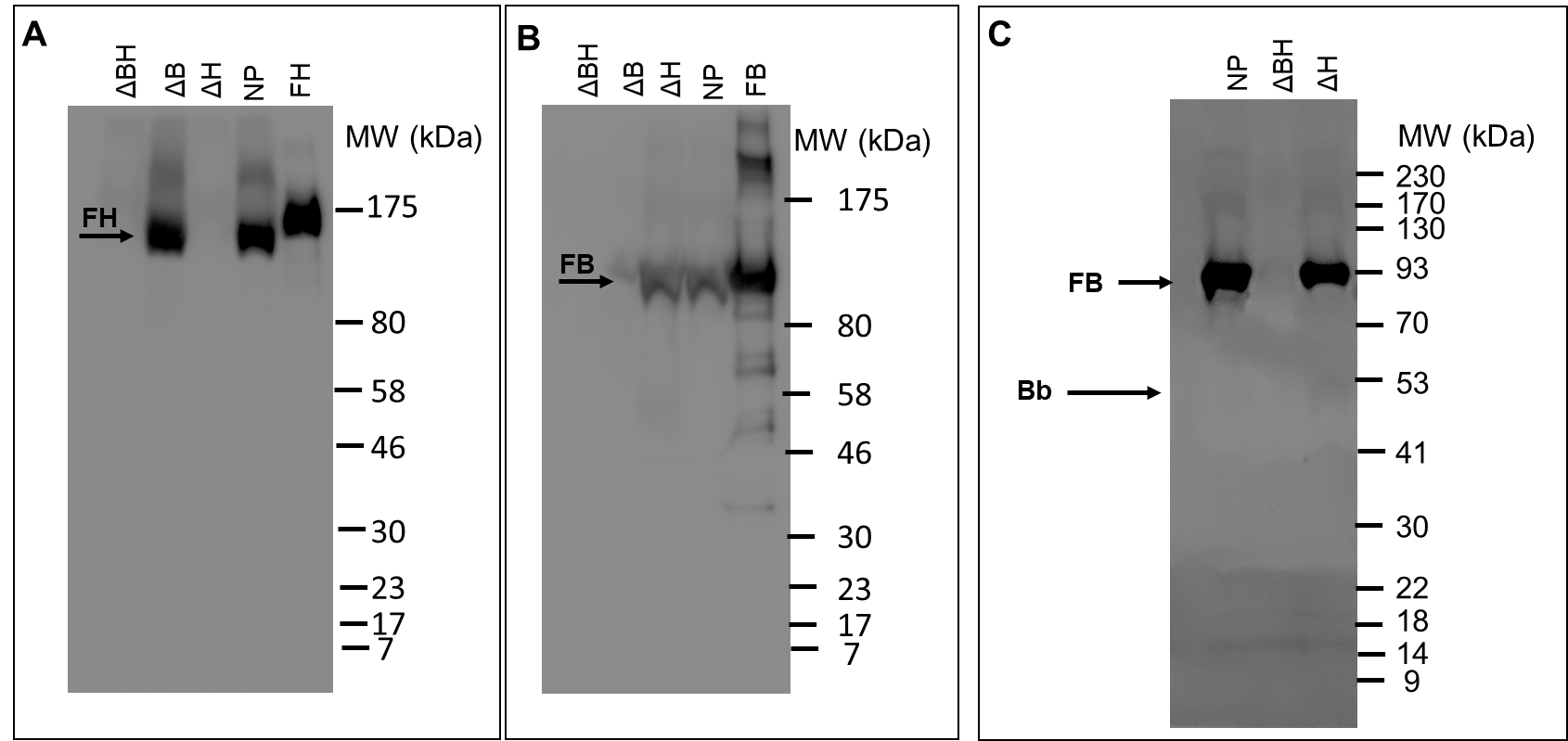

**Supplementary Figure 1. Western blot of FH and FB/FH affinity-depleted citrated plasma.** Factor H-depleted (ΔFH) and factor B/factor H-double depleted (ΔBH) plasmas alongside normal plasma (NP) and factor H (FH) and factor B (FB) protein control were run on SDS-PAGE under non-reducing conditions and bands visualised using monoclonal anti-FH 35H9) and anti-FB (JC1) antibodies. **(A)** FH depletion is confirmed in ΔH and ΔBH-depleted plasmas using anti-FH (35H9) mAb with FH protein as positive control. **(B)** FB depletion is confirmed in and ΔBH-depleted plasma using anti-FB (JC1) mAb with FB protein as positive control. (C) Western blot analyses determining intact FB and Bb fragment in citrated normal plasma (NP), alongside ΔH and ΔBH-double depleted plasma. FB and Bb is visualised by anti-FB/Bb (JC1) mAb. No residual complement alternative pathway activation (Bb generation) as a result of FH-affinity depletion was seen in citrated depleted plasma prior to analysis.

**Supplementary Figure 2**

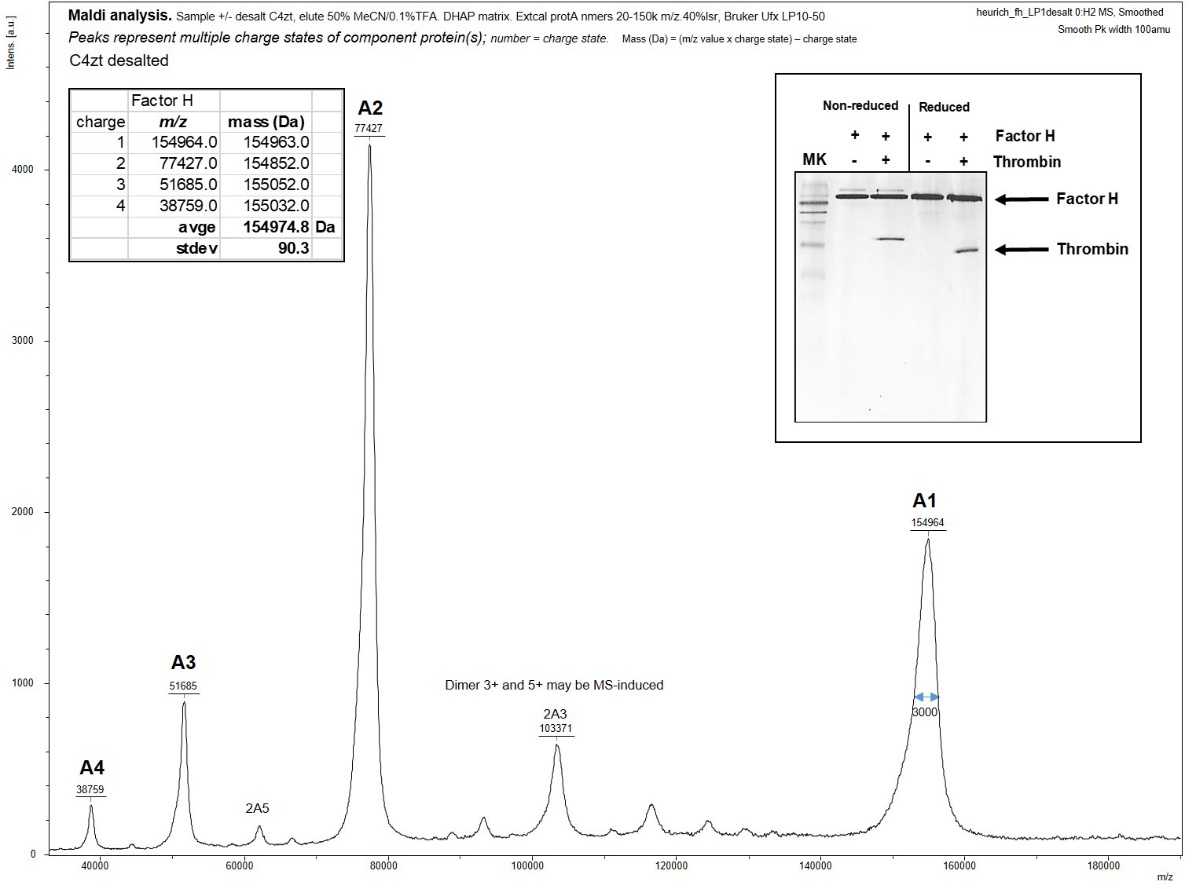

**Supplementary Figure 2. FH exposed to supraphysiological thrombin was not degraded and maintained its intact molecular weight. (A) T**hrombin (500nM) was incubated with 2μg FH for 1 hour at 37°C and protein bands resolved on SDS-PAGE (4-12% Tris-Glycine gradient gel) under non-reducing and reducing conditions alongside protein controls and protein bands were visualized by Coomassie stain. No FH cleavage fragments were observed by Coomassie. The FH band incubated with thrombin under non-reducing conditions was excised and subjected to in-gel MALDI analysis (sample +/- desalt C4zt, elute 50% MeCN/0.1%TFA. DHAP matrix. 20-150k m/z, Bruker Ufx LP10-50) to identify potential degradation of FH in the presence of thrombin. The most abundant mass was identified at 154975±90 Dalton confirming no significant mass loss and no FH proteolysis. Maldi-ISD analysis interrogated factor H N- terminal region and suggests that N-terminus is intact.
